# An ALS-associated mutation in human FUS reduces neurotransmission from *C. elegans* motor neurons to muscles

**DOI:** 10.1101/860536

**Authors:** Sebastian M. Markert, Michael Skoruppa, Bin Yu, Ben Mulcahy, Mei Zhen, Shangbang Gao, Michael Sendtner, Christian Stigloher

## Abstract

Amytrophic lateral sclerosis (ALS) is a neurodegenerative disorder that has been associated with multiple genetic lesions, including mutations in the gene FUS (Fused in Sarcoma), an RNA/DNA-binding protein. Expression of the ALS-associated human FUS in *C. elegans* results in mislocalization and aggregation of FUS outside the nucleus, and leads to impaired neuromuscular behaviors. However, the mechanisms by which mutant FUS disrupts neuronal health and function remain partially understood. Here we investigated the impact of ALS-associated FUS on motor neuron health using correlative light and electron microscopy, electron tomography, and electrophysiology. Expression of ALS-associated FUS impairs synaptic vesicle docking at neuromuscular junctions, and leads to the emergence of a population of large and electron-dense filament-filled endosomes. Electrophysiological recording of neuromuscular transmission revealed reduced transmission from motor neurons to muscles. Together, these results suggest a potential direct or indirect role of human FUS in the organization of synaptic vesicles, and reduced transmission from motor neurons to muscles.

**Summary statement:** An ALS-associated mutation in a trafficking protein disrupts the organization of the *C. elegans* neuromuscular junction.

## Introduction

Amytrophic lateral sclerosis (ALS) is a severe disease of the locomotor system where motor neurons progressively degenerate and muscles atrophy (Hardiman et al., 2017). ALS is hypothesized to be a synaptopathy (Fogarty, 2019), because the disease usually starts with dysfunction and degeneration of synapses, before axons and dendrites become dystrophic and the neurons undergo cell death (Chou, 1992). Most cases of ALS (∼90%) are considered ‘spontaneous’ – due to a combination of genetic and environmental factors. The remaining ∼10% of ALS cases are hereditary, and can be caused by mutation in several genes. A number of cellular defects have been implicated in the pathophysiology of ALS, including affected endosomal and receptor trafficking, as well as changes in autophagy and axonal transport (Burk and Pasterkamp, 2019).

Mutations in the gene FUS (fused in sarcoma) make up approximately ∼5% of hereditary ALS cases (Kwiatkowski et al., 2009; López-Erauskin et al., 2018; Vance et al., 2009). The FUS protein has a RNA recognition motif that contains a putative nuclear export signal, several disorganized domains, and a nuclear localization signal at the C-terminus (Kino et al., 2011; Lorenzo-Betancor et al., 2014; Murakami et al., 2012). Due to its nature as an RNA/DNA binding protein, FUS has been associated with a plethora of cellular functions, including translation, splicing, RNA transport, and as shown most recently, an involvement in DNA damage response signaling (Andersson et al., 2008; Lagier-Tourenne et al., 2010; Ratti and Buratti, 2016; Therrien et al., 2016; Wang et al., 2018). Under physiological conditions, FUS is predominantly located in the nucleus, but is able to shuttle between nucleus and cytoplasm in response to different stimuli (Gal et al., 2011; Vance et al., 2013). Pathological mutations are suggested to result in toxic gain-of-function, as ALS-associated FUS is prone to cytoplasmic accumulation. The potential role of nuclear loss of function through FUS sequestration in the cytoplasm is still being debated (An et al., 2019). Currently, the mechanisms by which mutant FUS leads to motor neuron degeneration are still largely enigmatic, with a dominating hypothesis that mutated FUS forms irreversible hydrogels that impair ribonucleoprotein (RNP) granule function (Murakami et al., 2015).

In a previous study, several variants of human ALS-associated FUS were ectopically expressed in the *C. elegans* nervous system (Murakami et al., 2012). These worms died prematurely, and had impaired motility, suggesting degeneration of the locomotor system through a dominant gain-of-function mechanism. The strength of the phenotype was correlated with the severity of human ALS caused by each variant, supporting ectopic FUS expression in the *C. elegans* nervous system as a promising approach with which to interrogate the mechanisms by which mutant FUS leads to disruption and degeneration of the nervous system. Here, we used behavioral, ultrastructural, and electrophysiological approaches to investigate how ALS-associated FUS impacts *C. elegans* neuromuscular transmission. We show that the organization of the neuromuscular junction is impacted by human FUS, and ALS-associated FUS impairs communication between motor neurons and muscles. These disease mechanisms may contribute to other forms of ALS, as well as other neurodegenerative diseases.

## Results

### Expression of ALS-associated FUS in the nervous system results in reduced lifespan and impaired motility

To determine the time window for comparative analysis, we first examined hermaphrodite worms that pan-neuronally express a pathological form of FUS, FUS C-terminal deletion (referred as FUS501 henceforth) for lifespan and motor function. Consistent with, and expanding previous results (Murakami et al., 2012, 2015), we observed a shortened lifespan of FUS501 animals (Figure 1A), and reduced motility (Figure 1B) compared to wild-type animals.

**Figure 1:**
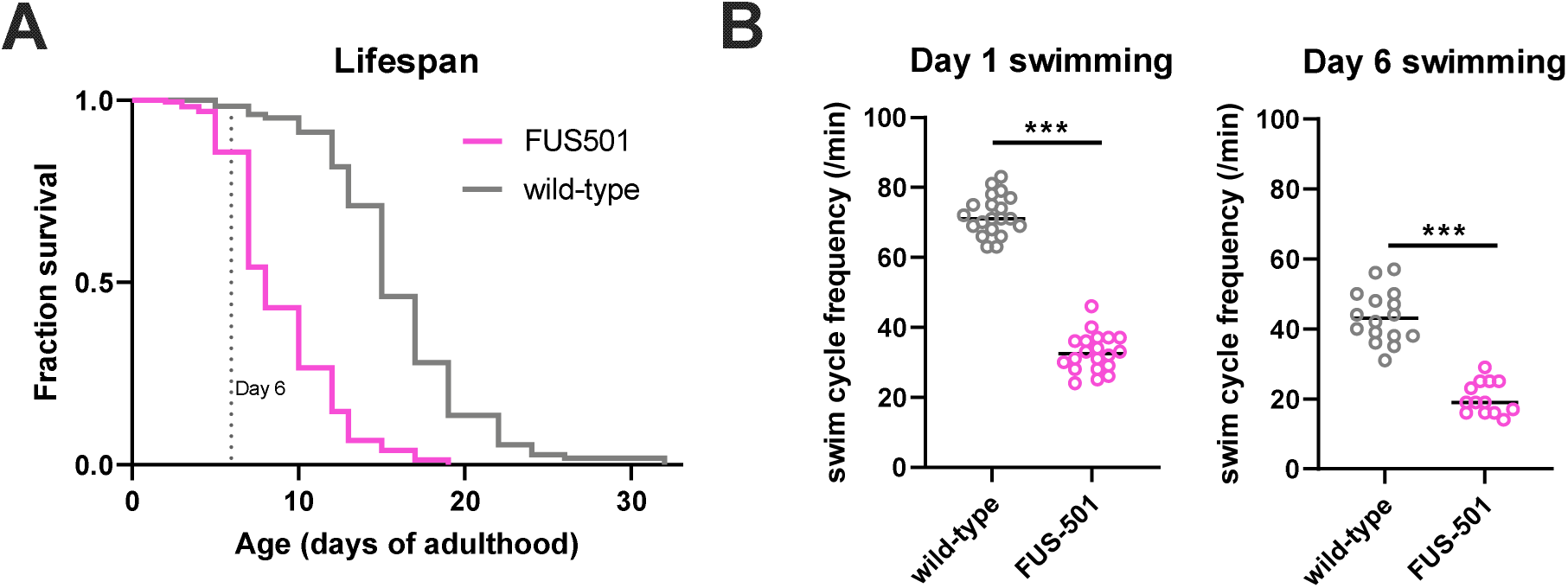
*C. elegans* expressing pan-neuronal human FUS501 exhibit lifespan and motility defects. (A) Animals expressing human FUS501 have a shorter median and maximum lifespan compared to wild-type animals (n≥94 deaths per genotype; log-rank test, p<0.0001). (B) *C. elegans* expressing FUS501 have defective swimming compared to wild-type (n≥10 per genotype; unpaired two-tailed Student’s t test, *** p<0.001).

Based on these results, we chose to proceed with comparative ultrastructural and functional analyses in FUS501 animals that were six days post-L4 stage adult. We reasoned that a pathological synaptic phenotype should already be pronounced prior to widespread degeneration, before causing secondary phenotypes that could make analysis more difficult. At this age, pathological features, including the FUS protein localization, aggregation, and motility were prominent, while animal survival was still high (Figure 1). As an additional control for structural analyses, we also included a line expressing wild-type FUS (FUSwt).

### Large endosomes with electron-dense inclusions are enriched at FUS501 neuromuscular junctions

In order to assess whether FUS501 motility defects result synaptic defects, we used electron tomography to map the organization of excitatory and inhibitory NMJs in wild-type (N2), as well as animals expressing wild-type human FUS (FUSwt) and FUS501.

N2 and FUSwt NMJs were composed of a presynaptic dense projection and associated pool of clear and dense core vesicles, and rarely contained large endosomes. However, large endosome-like organelles were enriched at the NMJs of animals expressing FUS501, and a population of large endosomes (>100nm) was exclusively present in FUS501 worms (Figures 2-4). This was accompanied by a reduction in the incidence of smaller (∼50 nm) diameter endosomes in FUS501 (Figure 4A). These A total of 196 electron tomograms of FUS501, FUSwt, or N2 synapses of NMJs in the ventral and dorsal nerve cords were reconstructed. Analyses were performed in a genotype-blinded manner. We selected 65 tomograms that contained active zones and were of sufficient quality for reliable vesicle analysis. Both GABAergic and cholinergic NMJs have been analyzed.

**Figure 2:**
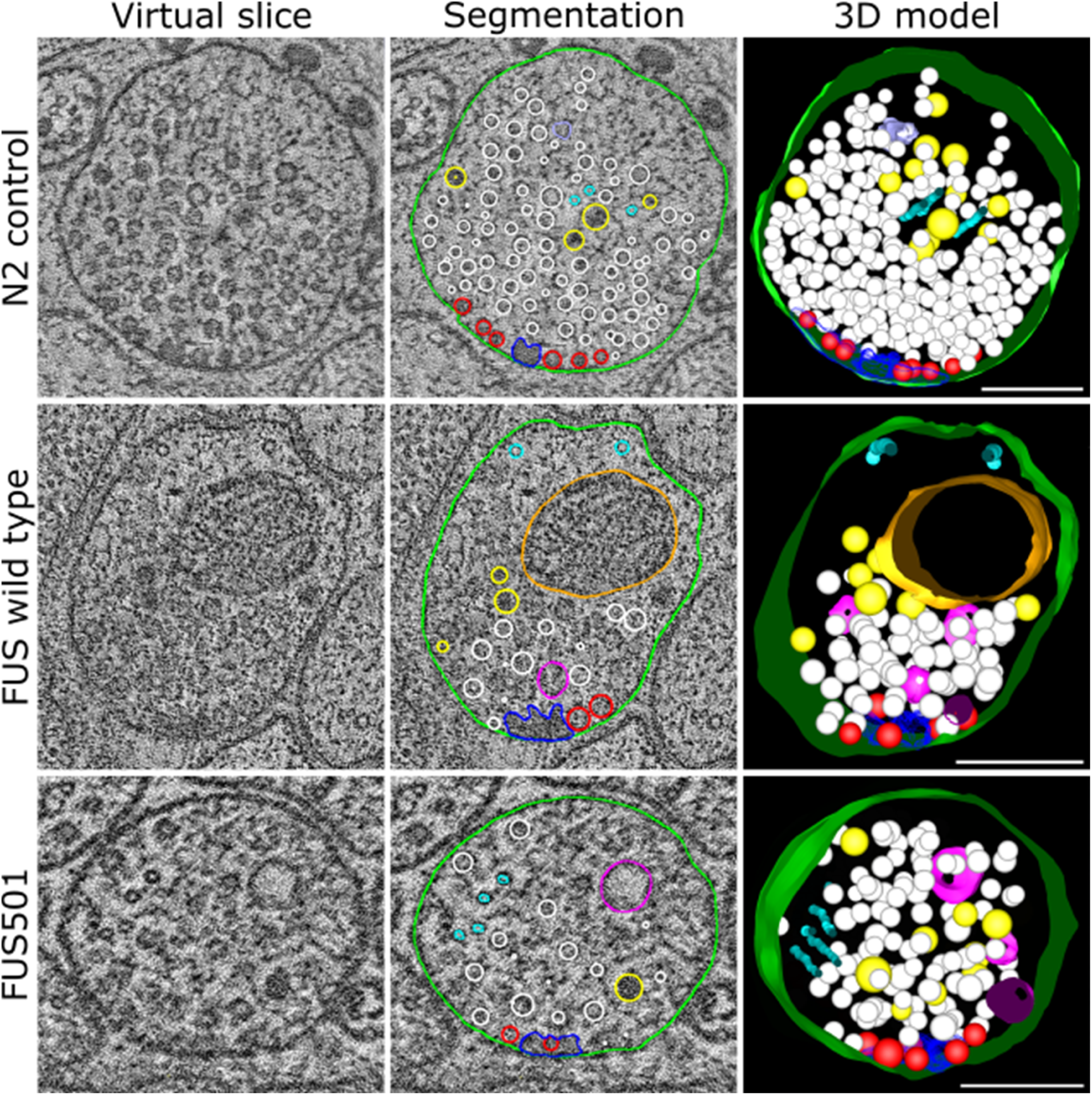
FUS501 affects the synaptic ultrastructure of cholinergic motor neurons. Shown are virtual ∼1nm slices from electron tomograms of cholinergic motor neurons, segmentations of these slices, and the 3D models of the whole tomograms. Segmented structures: plasma membrane (green), mitochondria (orange), dense projections (dark blue), microtubules (cyan), endoplasmic reticulum (lavender), dense core vesicles (yellow), clear core vesicles (white), docked clear core vesicles (red), and endosome-like structures (pink). Large, endosome-like structures appear in synapses affected by mutated FUS501, but not in FUSwt and N2 controls. FUSwt controls show smaller structures that presumably represent normal endosomes. Scale bars: 200 nm.

**Figure 3:**
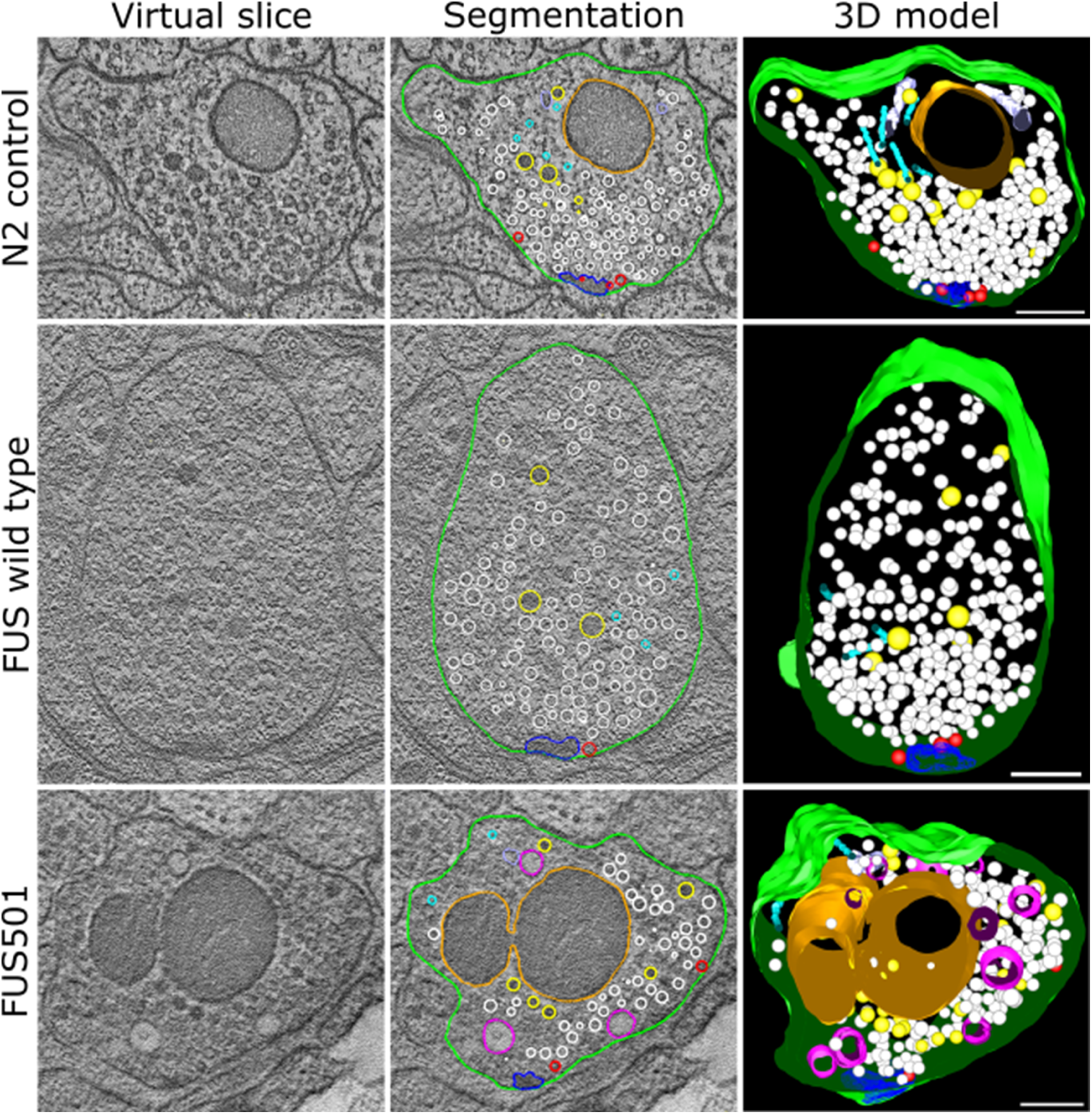
FUS501 affects the synaptic ultrastructure of GABAergic motor neurons. Shown are virtual ∼1nm slices from electron tomograms of GABAergic motor neurons, segmentations of these slices, and the 3D models of the whole tomograms. Segmented structures: plasma membrane (green), mitochondria (orange), dense projections (dark blue), microtubules (cyan), endoplasmic reticulum (lavender), dense core vesicles (yellow), clear core vesicles (white), docked clear core vesicles (red), and endosome-like structures (pink). Large, endosome-like structures appear in synapses affected by mutated FUS501, but not in FUSwt and N2 controls. Scale bars: 200 nm.

**Figure 4:**
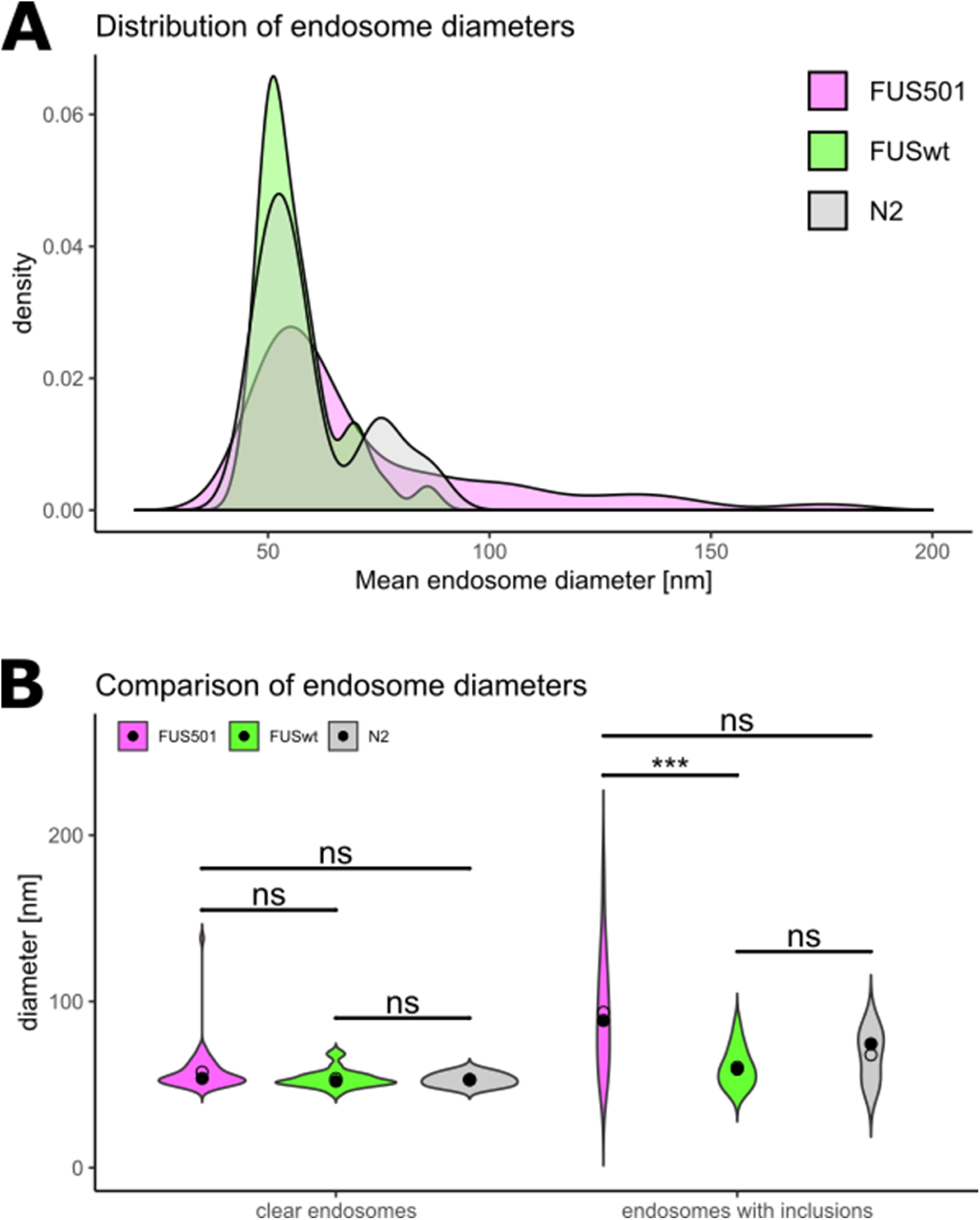
Larger endosomes that contain electron-dense filaments are caused by FUS501. Electron tomograms of 6 d hermaphrodite worms expressing FUS501 or FUSwt, or N2 worms were used to manually measure endosome diameters. For each endosome the average diameter calculated from the longest and shortest measured diameter was used for subsequent analysis. (A) Density plot of endosome diameters. FUS501 worms show populations of especially large endosomes not present in controls (arrows). (B) Comparison of endosome diameters in relation to the presence of electron-dense filaments. Statistical analysis via Mann-Whitney Wilcoxon test. Data are depicted as violin plots. Median (closed circles) and mean (open circles) are given on each plot.

We observed that NMJs of FUS501 worms showed enrichment in ultrastructural features only very rarely seen in both N2 and FUSwt controls. The most striking was the enrichment of large, mostly electron-light endosome-like organelles were mostly electron-light, and present in both cholinergic and GABAergic NMJs (Figures 2, 3). Most endosomes were spherical, although more complex networks also occurred (see Figure 5D).

**Figure 5:**
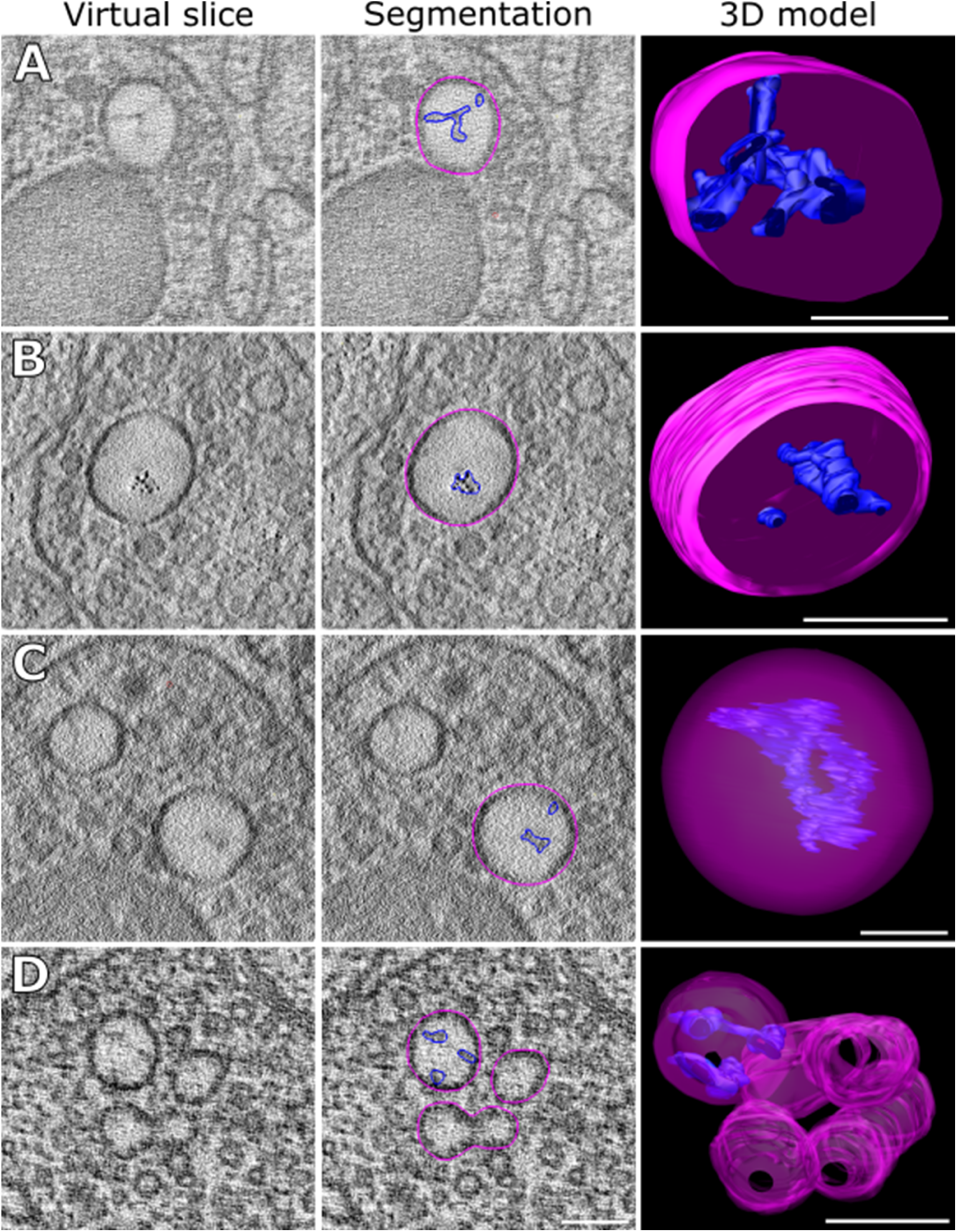
Morphology of the large endosomes. Examples of large endosomes (pink) in synapses of NMJs. Only endosomes in tomograms containing a dense projection are shown here. They appear large, with diameters of >80 nm, and often contain some electron-dense material (dark blue). (A) Typical example. The electron-dense content is often branched, as shown. (B) Electron-dense content appears partially in distinct dots. (C) Instance of a complete large endosome. (D) In one instance, large endosomes formed a group and “network” as shown. Scale bars: (A, B, D): 100 nm, (C): 50 nm.

In FUS501 mutants, approximately 41% of endosome-like structures had filamentous, electron-dense inclusions (35/85, from 26 tomograms). N2 and FUSwt animals also had a similar percentage of filled endosomes (36% and 41% respectively; 5/12 from 5 tomograms, and 14/39 from 17 tomograms). While ‘empty’ endosomes in each genotype were of a similar diameter, endosomes Figure 43). Overall, we found 50 clear endosomes and 35 filled ones in FUS501 (in 26 tomograms), 25 clear and 14 filled in FUSwt (in 17 tomograms), and 7 clear and 5 filled ones in N2 (in 5 tomograms). When comparing the diameters of the “clear” endosomes, there were no significant differences between FUS501 (53.8nm, MAD=4.8nm), FUSwt (52.0nm, MAD=4.4 nm), and N2 (53.0nm, MAD=5.2nm) (p-values: FUS501 vs. FUSwt: 0.089; FUS501 vs. N2: 0.24; FUSwt vs. N2: 1.0). However, the “filled” endosomes were larger in FUS501 worms containing electron-dense inclusions were larger than similar filled endosomes in N2 and FUSwt animals (Figure 4B) (88.5nm, MAD=35.6nm) compared to FUSwt (59.3nm, MAD=14.1 nm) and N2 (74.5nm, MAD=17.0nm).

The difference was highly significant for FUSwt (p-value=0.00074), but only a tendency for N2 (p-value= 0.098) (Figure 4). There was no significant difference between FUSwt and N2 (p-value=0.40). The low number of endosomes analyzed for N2 might explain the lack of statistical significance. Nevertheless, these data suggest that FUS501 expression in the *C.elegans* nervous system results in the emergence of a population of large endosome-like structures at the neuromuscular junction, through enlargement of endosomes that contain electron-dense filaments.

### Vesicle size at the neuromuscular junction is modified by human FUS

Given the presence of large endosomes in FUS501 terminals, we used automated reconstruction and classification tools (Kaltdorf et al., 2017, 2018) to reconstruct the vesicle pools from tomograms of cholinergic synapses. In total, we automatically reconstructed and classified 1,030 vesicles for FUS501, 164 vesicles for FUSwt, and 2,053 vesicles for N2 at the cholinergic NMJs. We found that clear synaptic vesicles (CCVs) had similar diameters between the three genotypes, although the small differences were statistically significant (median vesicle radii: 12.6 nm for FUS501, MAD=3.5 nm vs. 11.3 nm for FUSwt, MAD=4.4 nm vs. 13.2 nm for N2, MAD=3.5 nm; p-values: FUS501 vs. FUSwt: 0.0024; FUS501 vs. N2: 0.0049; FUSwt vs. N2: 8.9e-06) (Figure 6A). However, for dense core vesicles (DCVs), the diameter was similar in N2 and FUSwt N2 (median DCV radi: 23.5 nm for FUSwt, MAD=5.3 nm vs. 22.8 nm for N2, MAD=5.3 nm), but was significantly different in FUS501 (median DCV radius: 17.8 nm for FUS501, MAD=2.8 nm). DCVs made up 11.4 % of all vesicles in FUS501, 4.9 % in FUSwt, and 4.5 % in N2. Thus, in FUS501 worms, NMJ vesicle pools contain a higher proportion of DCVs that are smaller in size (Figure 6A).

**Figure 6:**
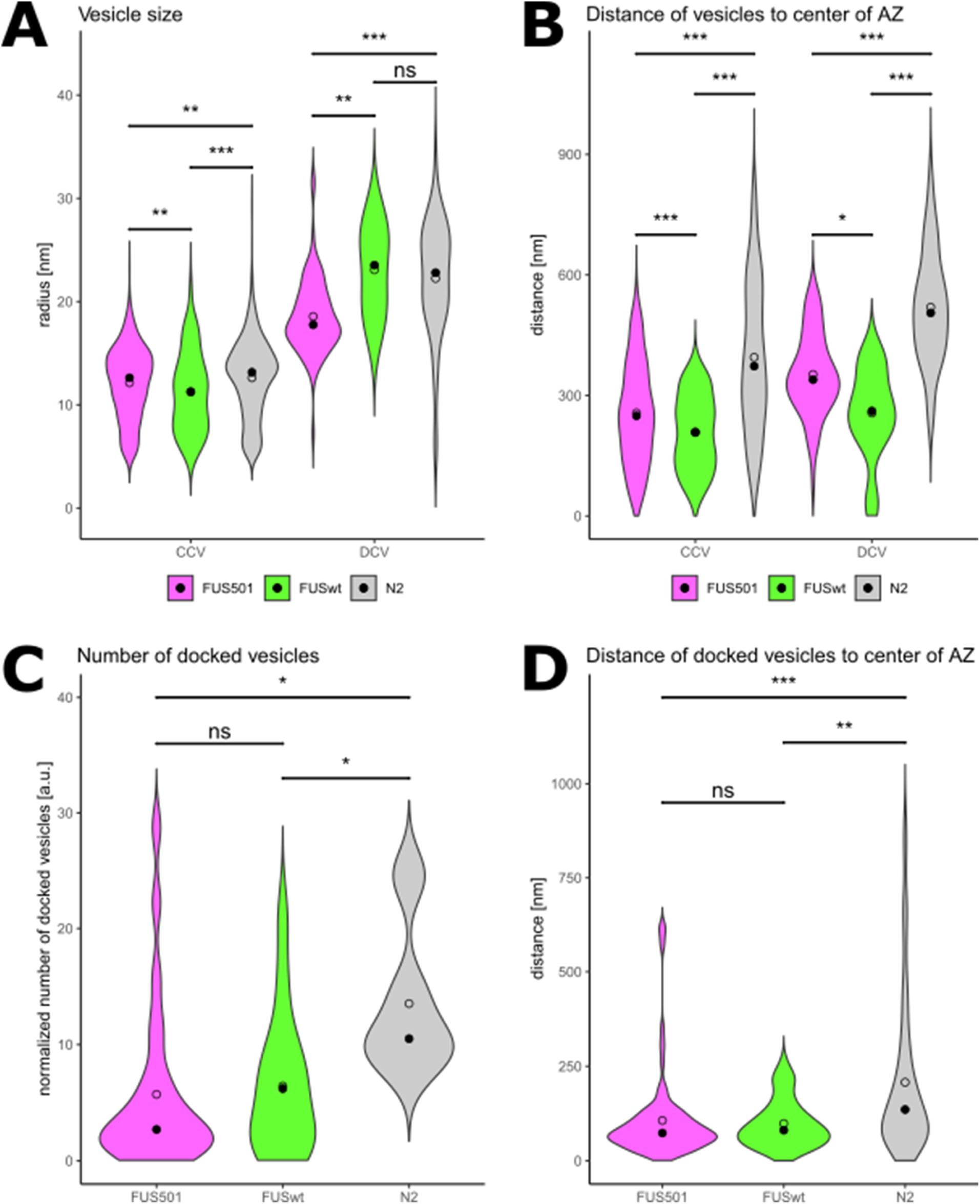
Expression of human FUS modifies the size, position and docking of vesicles at NMJs. Electron tomograms of 6 d hermaphrodite worms expressing FUS501 or FUSwt, or N2 worms were used for automated and manual analysis. Only cholinergic synapses were included. Statistical analysis via Mann-Whitney Wilcoxon test. Data are depicted as violin plots. Median (closed circles) and mean (open circles) are given on each plot. (A, B) Vesicle reconstruction and classification via the 3D ART VeSElecT and automated classification Fiji macros (Kaltdorf et al., 2017, 2018). (A) Linear distances of the center points of all vesicles to the center of the active zone (AZ) as given by the classification macro. (B) Vesicle radii as given by the classifiaction macro. (C, D) Analysis of vesicles docked to the plasma membrane via manual analysis. (C) Numbers of docked vesicles per tomogram normalized to approximate volume of the dense projection in a given tomogram. (D) Linear distances of docked vesicles to the center of the active zone (AZ).

### Human FUS disrupts the organization and docking of vesicles at the *C. elegans* neuromuscular junction

We further compared the distance between vesicles and the center of the presynaptic dense projection. In animals expressing human FUS (both FUSwt and FUS501) both clear- and dense-core vesicles were close to the presynaptic dense projection than N2 controls (Figure 6B). For both classes of vesicles, the effect of FUSwt was more pronounced than FUS501 (Figure 6B).

To assess how this altered distribution of vesicles may affect release properties at the NMJ, we quantified the number of docked vesicles at each synapse., but FUS501 showed also larger distances than FUSwt (249 nm for FUS501, MAD=141 nm vs. 208 nm for FUSwt, MAD=116 nm vs. 373 nm for N2, MAD=211 nm; p-values: FUS501 vs. FUSwt: 6.6e-05; FUS501 vs. N2: 2.2e-16; FUSwt vs. N2: 2.2e-16). For DCVs, distances for FUS501 and FUSwt differed significantly from each other (p-value=0.035), with greater distances for FUS501, but they both showed significantly smaller distances when compared to N2 (339 nm for FUS501, MAD=85.3 nm, p-value=4.5e-13 vs. 261 nm for FUSwt, MAD=80.7 nm, p-value=4.8e-05 vs. 505 nm for N2, MAD=127 nm) (Figure. 65B). These results show that vesicles are generally smaller in FUS501 and FUSwt, and located closer to the active zone.

To evaluate synaptic transmission, we analyzed docked vesicles in cholinergic NMJs (28 tomograms for FUS501, 16 for FUSwt, and 5 for N2). Both FUSwt and FUS501 had significantly fewer docked vesicles than N2 controls (number of docked vesicles per tomogram normalized to dense projection volume: 2.7 for FUS501, MAD=2.4 vs. 6.2 for FUSwt, MAD=5.9 vs. 10.5 for N2, MAD=3.4; p-values: FUS501 vs. FUSwt: 0.44; FUS501 vs. N2: 0.011; FUSwt vs. N2: 0.025) (Figure 6C). Docked vesicles in FUSwt and FUS501 were closer to the center of the presynaptic dense projection than N2 controls, similar to the global vesicle distribution described earlier When looking at the linear distances of docked vesicles to the center of the active zone, FUS501 and FUSwt showed no significant difference between each other, but again both were significantly smaller when compared to N2 (73.5 nm for FUS501, MAD=37.8 nm vs. 81.0 nm for FUSwt, MAD=54.9 nm vs. 136 nm for N2, MAD=102 nm; p-values: FUS501 vs. FUSwt: 0.47; FUS501 vs. N2: 0.00013; FUSwt vs. N2: 0.0034) (Figure 6D). Thus, in FUS501 and FUSwt fewer vesicles are docked, but those that are, are located closer to the active zone.

### FUS501 animals have reduced synaptic transmission from motor neurons to muscles

Given the defects in the structural organization of NMJ vesicle pools in animals expressing human FUS, we recorded the functional output of the NMJ by patch clamping muscle cells. The endogenous postsynaptic currents (mPSCs) reflect vesicle fusion events at the presynapse, that are due to either spontaneous release or endogenous activity in the motor neurons or upstream circuits, and subsequent detection of neurotransmitter (acetylcholine or GABA) by receptors on the muscle postsynapse (Richmond and Jorgensen, 1999). The frequency of mPSCs was no different from N2 controls in FUSwt, while FUS501 animals exhibited a >50% reduction in mPSC frequency (Figure 7A-B). There was no difference in mPSC amplitude between the three genotypes (Figure 7A, C). The presence of a frequency defect in the absence of amplitude defect suggests a reduction in presynaptic vesicle release from FUS501 motor neurons onto muscle cells. This could be due to defects in vesicle release at the level of individual NMJs, a reduction in the number of NMJs from motor neurons to muscle cells, or a combination of both.

**Figure 7:**
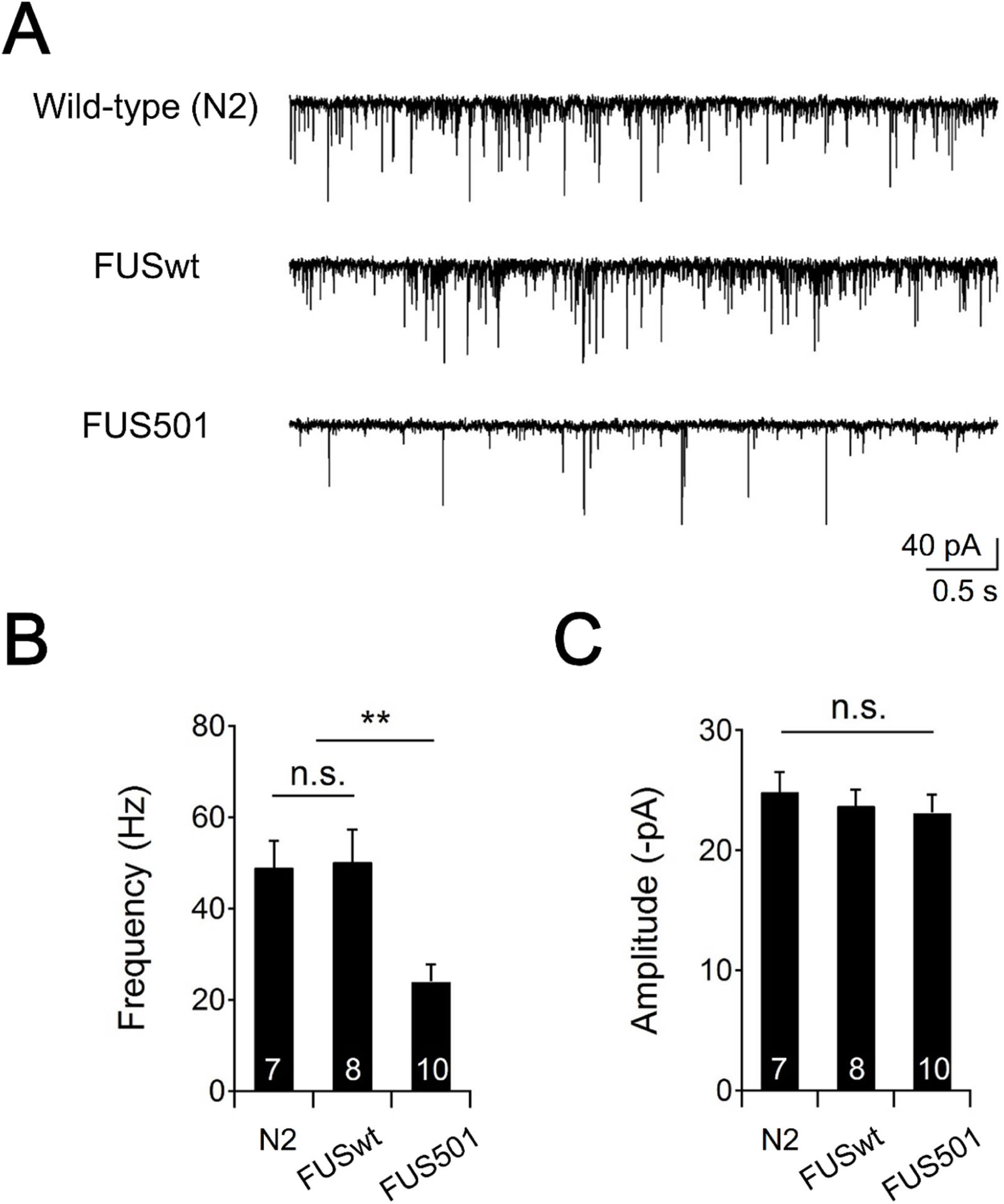
The frequency of minis is decreased in FUS501 transgenic animals. (A) Representative traces of spontaneous miniature synaptic currents in wild-type (N2), FUSwt, and FUS501 animals. Muscles were holding at −60 mV. (B) The frequency of minis was significantly decreased in FUS501 compared to wild-type (N2) or FUSwt animals. (C) The amplitude of minis showed no significant difference between strains. p-values: FUS501 vs. FUSwt: 0.0032; FUS501 vs. N2: 0.0018; FUSwt vs. N2: 0.89.

Overall, this suggest a defect in synaptic communication from motor neurons to muscles in FUS501 animals that is explained by reduced exocytosis at NMJs and/or a reduction in NMJ density. Because there was no change in mPSC amplitude, vesicle loading, and postsynaptic reception are likely unaffected at this timepoint (6 days of adulthood).

### FUS501 proteins aggregate in the nuclei and cytoplasm of motor neurons

Given that FUS501 NMJs harbor a pool of large, filament-filled endosome-like structures, we tested if the FUS501 protein was physically present at NMJs and could directly and locally be responsible for defects in neurotransmission in FUS501 animals.

Light microscopy studies have suggested that FUS501 forms aggregates in motor neuron cell bodies (Murakami et al., 2012, 2015; see also Figure S1). To localize FUS proteins in their ultrastructural context, we used super-resolution array tomography (srAT; Markert et al., 2017). FUS501 was located in aggregated clusters in the nucleus and the cytoplasm of motor neuron soma, but not in the nerve cords (Figure 8). The absence of detected FUS in motor neuron processes could be because it is not there, or because of a failure of detection due to insufficient preservation of small amounts of epitope. Nevertheless, FUS501 is not present in motor neuron processes at the high abundance seen in cell bodies. In our control worms, FUSwt staining was diffuse and limited to the nucleus (data not shown).

**Figure 8:**
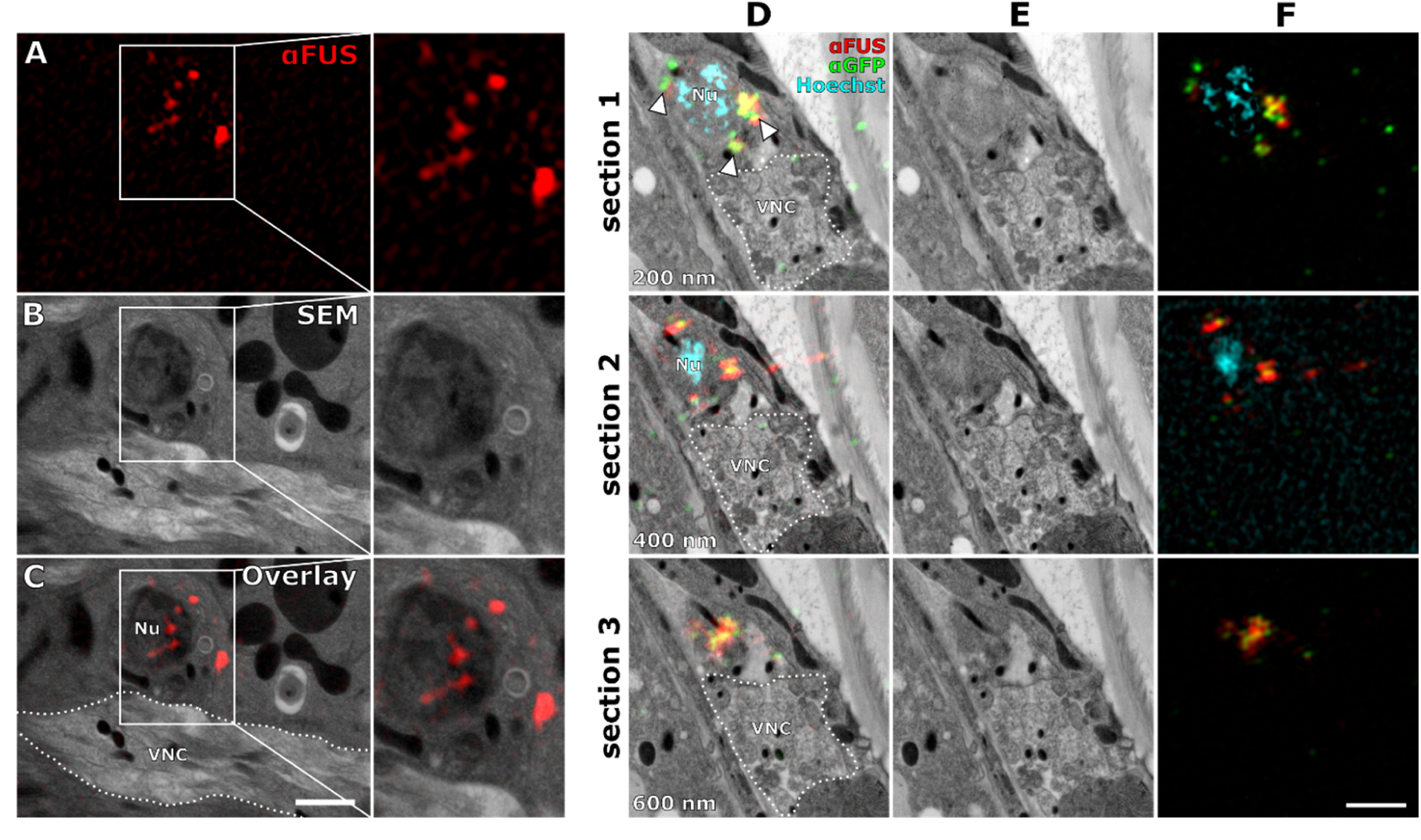
FUS501 is present in nucleus and cytoplasm of motor neurons but not in the nerve cords. (A) Immunofluorescence staining against mutated FUS acquired by SIM. (B) Scanning electron micrograph of the same region as in (A). (C) Overlay of (A) and (B) localizes FUS501 signals to the nucleus (Nu) and cytoplasm of a motor neuron, but does not show any signals on the synapses of the ventral nerve cord (VNC). (D) Scanning electron micrographs of the VNC region of an adult hermaphrodite worm expressing mutated FUS. Consecutive 200 nm sections were stained. Hoechst was used for correlation. FUS501 was stained via direct antibody (red) and via its GFP-tag (green). Both stainings overlap significantly (arrowheads). (E) Micrographs shown without overlay for reference. (F) Aligned SIM channels shown for reference. Scale bar: 1 µm.

## Discussion

In this study, we used ultrastructural and electrophysiological approaches to interrogate neuromuscular junctions of *C. elegans* expressing an ALS-associated variant of human FUS in the nervous system. The data suggest the resulting functional defect may be explained in part by changes in the organization of synaptic vesicle pools in the NMJ presynapse.

### FUS501-induced formation of large endosomal-like structures at NMJs

FUS501 worms had a population of unusually large vesicles at NMJs. The nature of these large intracellular vesicles remains to be better defined, but we consider them to be most likely endosomes: their morphology and location are consistent with endosomes described at *C. elegans* NMJs (i.e., they are roughly spherical and located within the synaptic vesicle pool; Watanabe, Liu, et al., 2013), and not other intracellular structures, such as autophagosomes (Meléndez et al., 2003), and the endoplasmic reticulum (ER). For example, autophagosomes feature double membranes, while our electron tomograms clearly show that they have single membranes (Figure 4). ER in our tomograms appeared as irregularly shaped tubes located at the synaptic periphery, whereas the putative endosomes appeared throughout synapses including regions close to the active zone. Thus, the population of large vesicles are most likely endosomes. Putative endosomes in each genotype, FUS501, as well as FUSwt and N2 control animals, were either ‘empty’ or contained electron-dense filamentous inclusions. The larger population of putative endosomes in FUS501 animals selectively contained inclusions. Interestingly, in Alzheimer’s disease and Down syndrome, endosomes have also been reported to be enlarged (Cataldo et al., 2008; Colacurcio et al., 2018), due to acceleration of endocytosis. However, our data suggest a decrease in endocytosis (see below). The electron-dense filamentous aggregates observed in these endosomes remain enigmatic. It is unknown if these filaments are native structures, of if the filaments in FUSwt or FUS501 contain FUS. It is also possible that such inclusions are a general aging phenotype, as they were also observed in age-matched (6 day adult) N2 controls. FUS501 aggregation may also promote aggregation of other proteins. It is well established that the *C. elegans* intracellular environment becomes more prone to aggregation across ageing, and this is exacerbated by the overexpression of aggregation-prone proteins (David et al., 2010; Huang et al., 2019; Walther et al., 2015).

Thus, FUS501 appears to selectively increase the diameter of endosome-like structures that contain inclusions. These endosome-like structures may be involved in bulk endocytosis, consistent with a possible defect in synaptic vesicle cycling in FUS501 nerve terminals.

### FUS501 interferes with neurotransmission, possibly in part through disrupting synaptic vesicle organization

Expression of human FUS, whether FUSwt or FUS501, resulted in changes to vesicle size and distribution at the neuromuscular junction. It also resulted in fewer docked vesicles. Though this was observed at both FUSwt and FUS501 terminals, functional defects in synaptic transmission measured by electrophysiology were only observed at FUS501 terminals. It is possible that reduced ePSCs in FUS501 was due to 1) reduced vesicle release from individual NMJs; 2) reduced number of NMJs innervating each muscle cell; or 3) a combination of both.

If changes in vesicle organization and docking may contribute to defects in neurotransmission at FUS501 NMJs, additional factors are likely at play. These may be within the synaptic terminal (e.g. changes in coupling of synaptic vesicles to voltage-gated calcium channels; Chang and Martin, 2016), within the motor neuron (e.g. reduced excitability; Guo et al., 2017; Liu et al., 2016; Naujock et al., 2016), or in upstream circuits that modulate endogenous motor neuron activity.

In ALS, excitotoxicity is a topic of concern with regards to neuron degeneration (Fogarty, 2019). It is thought that neurons are driven to decay by hyper-excitability for several forms of ALS (Fogarty, 2018). However, for FUS-mediated mouse model, motor neuron degeneration was reported to be preceded by hypo-excitability (Martínez-Silva et al., 2018), echoing the high heterogeneity of ALS (Hardiman et al., 2017). Our results support the hypo-excitability model and are thus consistent with the conclusion from the mouse FUS models (Kong et al., 2009; Martínez-Silva et al., 2018; Ruiz et al., 2010).

For both ultrastructural and functional analyses, we used two lines as controls in all experiments, N2 and FUSwt. While wild-type human FUS clearly has mild effects the ultrastructure of its nervous system, such effects did not cause significant reduction in synaptic transmission.

### srAT reveals the ultrastructural context of FUS localization

Consistent with the implication of fluorescent microscopy studies (Murakami et al., 2012, 2015) we observed clusters of FUS in the nucleus and cytoplasm with srAT (Figures 8, S1). While srAT allowed us to localize FUS with high precision and accuracy, we did not find FUS accumulations in axons, most likely due to the low abundance of preserved epitopes, and insufficient sampling of our serial sections. A previous study (Murakami et al., 2015) showed strong evidence that irreversible FUS hydrogels are associated with RNP granules, which have been described by electron microscopy using samples preserved by a different protocol (Biggiogera et al., 1997; Jokhi et al., 2013; Souquere et al., 2009). It may be possible to visualize ultrastructural features of FUS aggregates or the RNP granules they attach to using different preservation protocols.

### A reduction of protein translation might account for endosome and vesicle docking defects

FUS has many functions related to DNA and RNA processing and maintenance (Ratti and Buratti, 2016), and knock-out leads to perinatal death in mice (Hicks et al., 2000). Certain mutation alleles, like FUS501, cause degeneration of motor neurons via a dominant gain-of-function effect (Murakami et al., 2012). Irreversible hydrogels formed by mutated FUS impair RNP granule function. This reduces the rate of new protein synthesis (Murakami et al., 2015). This has a systemic effect on neurons, however, the effect on axons and synapses is likely particularly detrimental, since such intracellular compartments have been shown to be heavily dependent on local translation regulation by RNP granules in mouse (Akbalik and Schuman, 2014; Holt and Schuman, 2013; Jung et al., 2014) and in *C. elegans* (Yan et al., 2009). Furthermore, the replacement of murine FUS with a mutated human form activates an integrated stress response and inhibit local intra-axonal protein synthesis in hippocampal neurons and sciatic nerves, resulting in synaptic dysfunction (López-Erauskin et al., 2018). A recent study found that transcription of an acetylcholine receptor is compromised in FUS-mediated ALS, thus supporting the idea that FUS affects gene expression (Picchiarelli et al., 2019).

It is thus plausible that the ultrastructural defects that we found is caused by a reduced protein synthesis. Under physiological conditions, vesicles are recycled via the ultrafast endocytosis pathway (Watanabe, Liu, et al., 2013; Watanabe, Rost, et al., 2013). After endocytosis, vesicles are regenerated in a clathrin-dependent manner (Watanabe et al., 2014). Shortage of clathrin or other components of this pathway could cause an accumulation of endocytosed membrane represented by large endosomes. This hypothesis is supported by our result that median distance of vesicles to the active zone is reduced in FUS worms, which is consistent with a reduction of vesicle pool size.

## Materials and methods

### Worm strains

All *C. elegans* worms were maintained according to standard methods (Brenner, 1974). Three strains used in this study are ZM9566, which ectopically and panneuronally expresses FUS501, ZM5838, which ectopically and panneuronally expresses wild-type FUS, both under the control of the *Prgef-1* promoter, and ZM9569, which was used as the N2 control. ZM9566 and ZM5838 were derived by outcrossing against N2 Bristol 6 times to remove any potential background accumulation (Murakami et al., 2012). ZM9569 was the non-transgenic progenies recovered from the same outcross that derived ZM9566.

### Lifespan

*C. elegans* were synchronized at the L4 stage and manually picked to fresh plates every 1-2 days to prevent contamination by progeny. Animals that were immotile and did not respond to gentle prodding with a platinum wire were scored as dead. Animals that bagged (died due to internal hatching of progeny) or crawled up the walls of the plate and desiccated were censored from the lifespan analysis. The strains used in the lifespan experiment were ZM9566 (panneuronal FUS501 expression) and ZM9569 (wild-type).

### Thrashing assay

In thrashing experiments, animals cultured 16 h post L4 larva were scored as Day 1 adults. Individual animals were transferred into 5 μl of M9 buffer on a glass slide. Thirty seconds after the transfer, the frequency of the body bending was counted for 1 min. A single thrashing was defined as a complete sinusoidal movement through the head and tail. The same protocol was used to assess the motor function in Day 6 adults.

### High-pressure freezing

The samples are subjected to >2100 bars of pressure and cooling rates of >20,000 K/s. All samples used in this study were cryo-immobilized using an EM HPM100 (Leica Microsystems) high-pressure freezing machine. The procedure was to use freezing platelets (Leica Microsystems) with 100 µm recesses. They were slightly overfilled with OP50 paste (see below) and then worms were transferred into the platelet. A second platelet with a flat surface was placed on top as a lid. The samples were processed and then stored in liquid nitrogen until freeze-substitution.

### *E. coli* OP50 bacteria paste

A 100 ml volume of *E. coli* OP50 overnight culture was pelleted at 1,500 ×g, washed with 400 µl 20% bovine serum albumin (BSA) in M9 (Stiernagle, 2006) (3.0 g KH_2_PO_4_, 6.0 g Na_2_HPO_4_, 0.5 g NaCl, 1 ml 1 M MgSO_4_, H_2_O to 1 l; sterilize by autoclaving), spun down again, and carefully re-suspended in 20 µl 20% BSA in M9.

### Freeze-substitution and resin embedding optimized for structure preservation

The protocol is originally based on (Weimer, 2006). A description of the individual steps of the freeze-substitution and resin embedding can be found in (Stigloher et al., 2011).

### Freeze-substitution and resin embedding for retention of antigenicity

Again, the protocol is originally based on (Weimer, 2006). We published a detailed description of the individual steps of the freeze-substitution and resin embedding previously (Markert et al., 2017).

The protocol uses potassium permanganate alone as fixative. After substitution, samples are embedded in LR White resin (Medium Grade Acrylic Resin, London Resin Company Ltd.). This protocol was used for all srAT applications.

### Ultramicrotomy for srAT

We published a detailed description of the individual steps previously (Markert et al., 2017).

Briefly, 100 nm sections were produced using a special diamond knife with a boat large enough to accommodate glass slides (histo Jumbo diamond knife, DiATOME). Slides were submerged in the boat before sectioning. Then the desired number of sections was cut without interruption. If necessary, a long ribbon was carefully divided into smaller ribbons using two mounted eyelashes.

### Ultramicrotomy for electron tomography

For imaging with a 200 kV TEM, we used sections up to 250 nm with good results. Sections between 150 and 200 nm in thickness were favored.

#### Immunostaining

Ultrathin sections of LR White-embedded tissue were immunostained for srAT. Ultramicrotomy exposed epitopes at the section surface. Thus, sections could be stained, even though antibodies do generally not penetrate the resin.

The detailed protocol has been published before (Markert et al., 2017). In brief, sections were placed in a humid chamber, and blocking buffer was applied to the sections prior to staining with primary and secondary antibodies. They were washed with buffer and then stained with a DNA stain, where applicable. Lastly, sections were mounted with Mowiol and stored at 4*^◦^*C for up to a week before fluorescence imaging.

#### Imaging for srAT

The following workflow of srAT imaging has been published in detail (Markert et al., 2016, 2017).

#### Contrasting and carbon coating

Contrasting was achieved by incubation of the sections in 2.5% uranyl acetate in ethanol for 15 min and 50% Reynolds’ lead citrate (Reynolds, 1963) in ddH_2_O for 10 min. While incubating in lead citrate, sodium hydroxide pellets were placed around the samples to decrease local carbon dioxide concentration, as carbon dioxide forms a precipitate with lead citrate. In between contrasting steps, the slides were washed first in ethanol, then in 50% ethanol in ddH_2_O, and finally in ddH_2_O. After contrasting, they were thoroughly washed in ddH_2_O and dried with pressurized air.

Carbon coating was usually performed right after contrasting. Based on the shadow cast on a piece of white filter paper, we estimated the thickness of the carbon layer necessary to quench charging to be about 10 nm. If in doubt, a thin carbon layer was tested first. In case there was charging, the samples were coated again until it ceased. If the carbon layer was much too thick, SEM imaging resolution was negatively affected.

#### Scanning electron microscopy (SEM)

All scanning electron microscopy was performed with a field emission scanning electron microscope JSM-7500F (JEOL, Japan) with LABE detector (for back scattered electron imaging at extremely low acceleration voltages). The acceleration voltage was kept at 5 kV at all times, the probe current was kept at 0.3 nA. Imaging was generally performed at 10 µA beam current or lower. The resulting working distance was 6-8 nm.

#### Correlation

Correlation strategies and procedures were described in detail previously (Markert et al., 2017). Briefly, there were two main strategies used: manual and semi-automatic. Manual correlation was performed with Inkscape (version 0.91; http://www.inkscape.org). For semi-automatic correlation, the eC-CLEM plugin (Paul-Gilloteaux et al., 2017) for the software Icy (de Chaumont et al., 2012) was used. Here, a small number of markers are placed on structures visible in both imaging modalities. The images are then superimposed automatically based on these manually placed markers.

### Preparation of sections and imaging for electron tomography

Imaging was performed with the SerialEM (Mastronarde, 2005) and IMOD (Kremer et al., 1996) software packages. A 200 kV JEM-2100 (JEOL) electron microscope equipped with a TemCam F416 4k×4k camera (Tietz Video and Imaging Processing Systems) was used for all TEM imaging and electron tomography.

#### Contrasting

Contrasting was achieved by floating the grids sections down on drops of 2.5% uranyl acetate in ethanol for 15 min and 50% Reynolds’ lead citrate (Reynolds, 1963) in ddH_2_O for 10 min. During incubation, samples were covered to minimize evaporation. During incubation in lead citrate, sodium hydroxide pellets were placed around the samples to decrease local carbon dioxide concentration. Carbon dioxide forms precipitates with lead citrate. In between contrasting steps, the grids were washed first in ethanol, then in 50% ethanol in ddH_2_O, and finally in ddH_2_O. After contrasting, they were thoroughly washed in ddH_2_O and blotted dry with filter paper.

#### Carbon coating and placement of gold fiducials

Grids used for electron tomography were coated with a thin layer of carbon to prevent charging during imaging at high tilt angles. The carbon layer had an approximate thickness of three nm.

Gold fiducials were used to facilitate tomogram reconstruction. To achieve fiducial placement, a non-specific antibody conjugated with 10 nm gold particles was used. The antibody was diluted 1:10 with ddH_2_O and 50 µl of this dilution were pipetted on a piece of clean parafilm. The carbon-coated grids were then floated on the drop for 10 min on each side, with a single wash in ddH_2_O in between and at the end. A single wash meant that the grid was submerged in water for one second and then immediately dried with a filter paper. The gold fiducial placement was always performed right after carbon coating or at most a few hours after. For unknown reasons, longer delays caused very pronounced electron-dense precipitation on the sections, making them unsuitable for imaging in extreme cases.

#### Acquisition of tilt series

Tilt series for this thesis were acquired either from 60° to −60° or from 70° to −70°. Double tilts were performed where appropriate and possible, i.e., tilt series from a region of interest were acquired in two orthogonal tilt axes. This was achieved by manually rotating the grid by about 90° in the sample holder. Double tilts improved tomogram quality significantly. They were not performed when the tomogram of a single axis was sufficient to answer the specific questions.

#### Tomogram reconstruction

All tomograms were reconstructed with the eTomo software from the IMOD package (Kremer et al., 1996). Gold fiducials were always included to improve the alignment of the tilt series. For the step of tomogram positioning, the option “find boundary model automatically” was used. Manual adjustments of the boundary model were almost never necessary. Tomograms were always created using the “Back Projection” algorithm.

#### Segmentation and 3D reconstruction

Segmentation and 3D reconstruction were performed with the 3dmod software from the IMOD package (Kremer et al., 1996). All structures except for the vesicles were segmented as closed objects using the “sculpt” tool. Clear core and dense core vesicles were annotated as perfect spheres by creating a point in the center of a vesicle using the “normal” drawing tool. This point was then resized with the mouse wheel to match the outer dimensions of the given vesicle. Global quality of points was set to 4 to achieve smooth spheres and the “drawing style” of points was set to “fill” to obtain closed surfaces. All other objects except the dense projections were meshed to obtain closed surfaces here as well. Dense projections were left with the default drawing style “lines”. The “interpolator” tool was used whenever appropriate. For large structures like the plasma membranes, gaps of 20 virtual sections or more were linearly interpolated. For mitochondria and microtubules typically gaps of 10 sections were interpolated. Larger spherical structures like endosomes were interpolated with the “spherical” option. Dense projections were not interpolated.

#### Quantitative analyses

Automatic vesicle reconstruction from electron tomograms was performed via macros for the open source image processing software Fiji (Schindelin et al., 2012) as described in (Kaltdorf et al., 2017). They were then automatically classified into clear core and dense core vesicles according to (Kaltdorf et al., 2018). Manual adjustments of the outcomes were not performed. However, if overall classification results for a given tomogram were not satisfactory, this tomogram was excluded from analysis. The active zone was determined manually for the classification macro. A point on the plasma membrane that is closest to the center of gravity of the dense projection seen in a given tomogram was set as the center of the active zone. The center of gravity of the dense projection was chosen by visual judgment of the user during the macro workflow. Manual vesicle reconstruction from electron tomograms was performed via 3dmod from the software package IMOD (Kremer et al., 1996). The center points of vesicles were set by the user’s judgment and set as centers of spheres with the approximate outer diameter of the vesicles.

The dense projections were segmented manually, and their center of gravity was determined with the “imodinfo” function of IMOD. The center of the active zone was defined as the intersection of the inner plasma membrane and an orthogonal line through the center of gravity of the dense projection. Linear distances of vesicles to the active zone were measured with the “measure” tool in 3dmod from the centers of the vesicles to the center of the active zone.

Average endosome diameters were calculated from the longest and shortest diameter of a given endosome measured manually on the virtual tomogram slice where the endosome appeared largest.

#### Statistical analyses

Statistical analyses and their representations were performed with R (R Core Team 2019). The Mann-Whitney-Wilcoxon test was used to determine statistical significance. The following significance levels were applied: * p<0.05, ** p<0.01, *** p<0.001.

Variability of quantitative data samples was measured via median absolute deviation (MAD). The MAD is more robust against outliers and suitable for non-parametric data, i.e., data that does not show normal distribution (Pham-Gia and Hung, 2001). It is defined as the median of the absolute deviations of the data’s median:

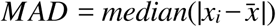

for a univariate data-set *x*_1_*, x*_2_*, …, x_n_* where 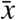 is the median of the data: 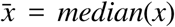.

#### Electrophysiology

The dissection of the *C. elegans* was described previously (Richmond and Jorgensen, 1999). Briefly, 6d hermaphrodite adults were glued to a sylgard-coated cover glass covered with bath solution. The integrity of the ventral body muscle and the ventral nerve cord were visually examined via DIC microscopy, and muscle cells were patched using fire-polished 4-6 MΩ resistant borosilicate pipettes (World Precision Instruments, USA). Membrane currents were recorded in the whole-cell configuration by a Digidata 1440A and a MultiClamp 700A amplifier, using the Clampex 10 software and processed with Clampfit 10 (Axon Instruments, Molecular Devices, USA). Data were digitized at 10-20 kHz and filtered at 2.6 kHz. The recording solutions were as described in our previous studies (Gao and Zhen, 2011). Specifically, the pipette solution contains (in mM): K-gluconate 115; KCl 25; CaCl_2_ 0.1; MgCl_2_ 5; BAPTA 1; HEPES 10; Na_2_ATP 5; Na_2_GTP 0.5; cAMP 0.5; cGMP 0.5, pH7.2 with KOH, ∼320 mOsm. The bath solution consists of (in mM): NaCl 150; KCl 5; CaCl_2_ 5; MgCl_2_ 1; glucose 10; sucrose 5; HEPES 15, pH7.3 with NaOH, ∼330 mOsm. Leak currents were not subtracted. All chemicals were from Sigma. Experiments were performed at room temperatures (20-22°C)

## Acknowledgements

We like to thank Philip Kollmannsberger for helpful discussions. SMM was supported by the Studienstiftung des Deutschen Volkes.

---

**Figure S1:**
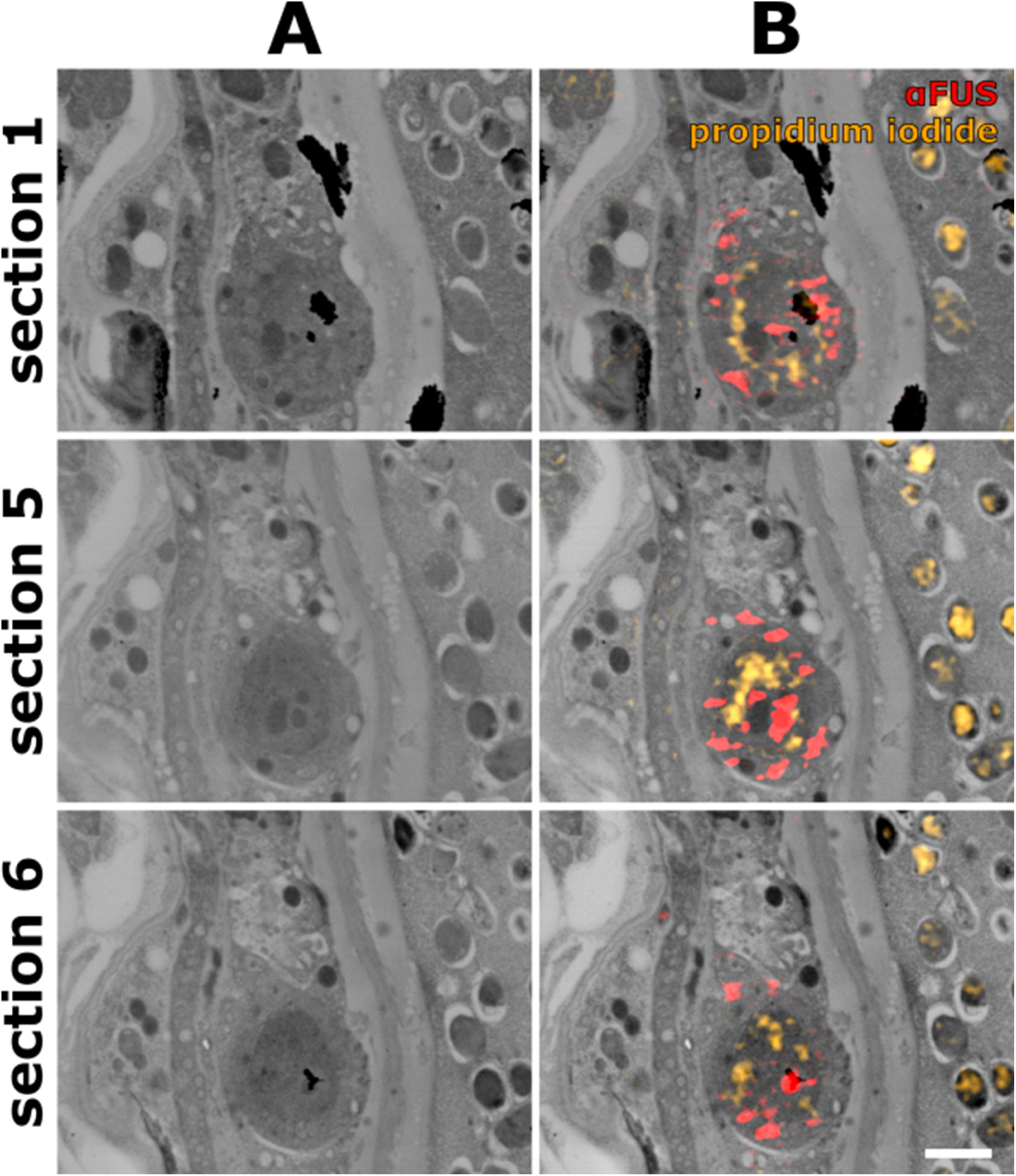
A selection of sections through a VNC consistently shows localization of mutated FUS in both cytoplasm and nucleus. (A) Scanning electron micrographs of the VNC region of an adult hermaphrodite worm expressing mutated FUS. (B) The images of (A) overlaid with their corresponding fluorescent SIM images. Propidium iodide was used for correlation. FUS501 signals are consistently localized to both the nucleus and the cytoplasm of a motor neuron. The patchy staining suggests the presence of FUS accumulations. Scale bar: 1 µm.

**Figure S2:**
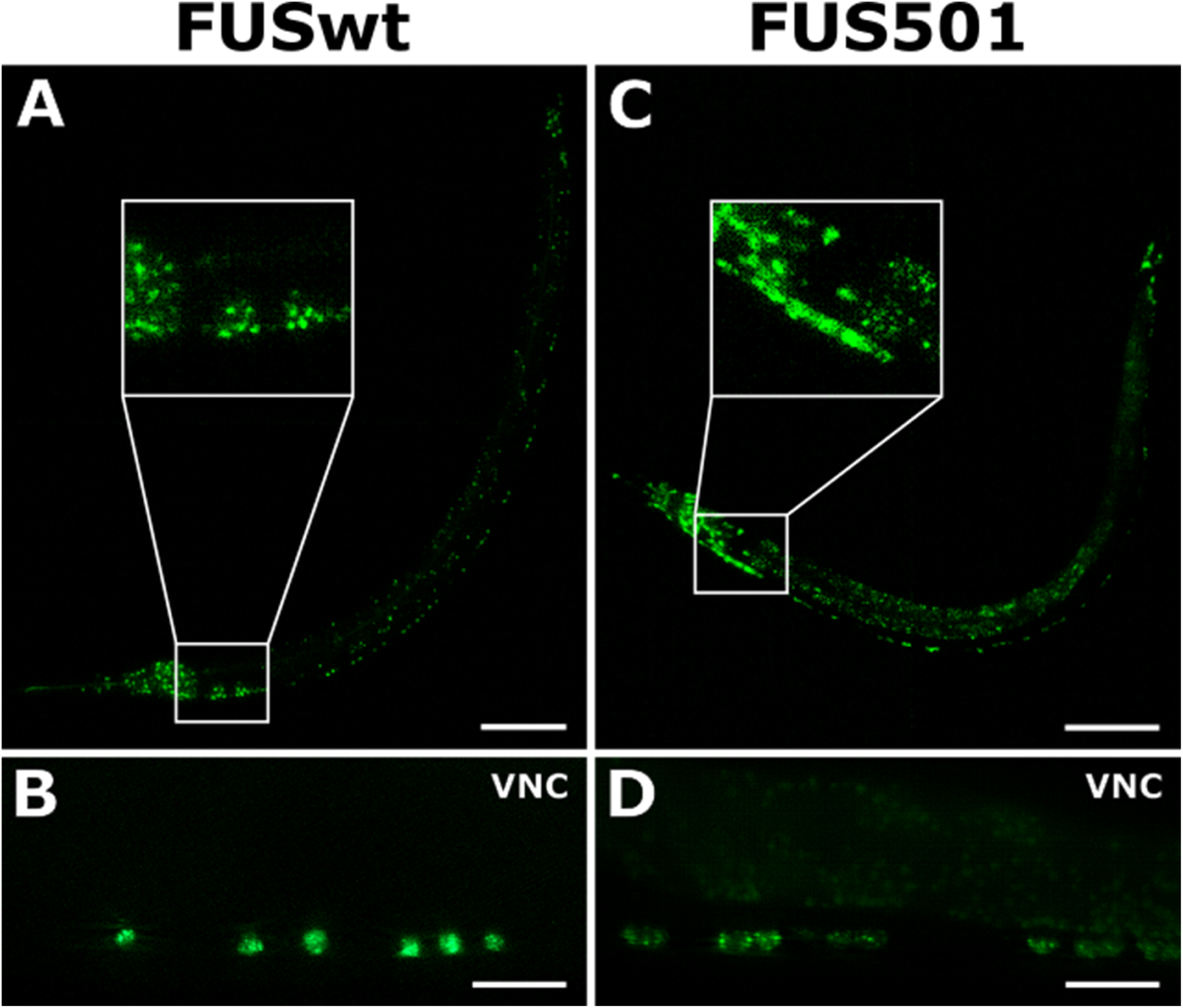
Structured illumination micrographs of FUSwt and FUS501 expression in living worms. FUSwt and FUS501 were imaged in living L3 larvae via their GFP-tags. Shown are maximum intensity projections of z-stacks. (A) Overview of FUSwt expression in the whole larva. Scale bar: 50 µm. (B) FUSwt expression in the ventral nerve cord (VNC). Scale bar: 10 µm. Note how FUSwt is located in distinct circular patterns, presumably within nuclei. (C) Overview of FUS501 expression in the whole larva. Scale bar: 50 µm. (D) FUS501 expression in the ventral nerve cord (VNC). Scale bar: 10 mm. Note how FUS501 is distributed more diffusely and forms small puncta, presumably aggregates.

## References

Akbalik, G. and Schuman, E.M. (2014), “mRNA, Live and Unmasked”, Science, Vol. 343 No. 6169, pp. 375–376.

An, H., Skelt, L., Notaro, A., Highley, J.R., Fox, A.H., La Bella, V., Buchman, V.L., et al. (2019), “ALS-linked FUS mutations confer loss and gain of function in the nucleus by promoting excessive formation of dysfunctional paraspeckles”, Acta Neuropathologica Communications, Vol. 7 No. 1, p. 7.

Andersson, M.K., Ståhlberg, A., Arvidsson, Y., Olofsson, A., Semb, H., Stenman, G., Nilsson, O., et al. (2008), “The multifunctional FUS, EWS and TAF15 proto-oncoproteins show cell type-specific expression patterns and involvement in cell spreading and stress response”, BMC Cell Biology, Vol. 9, p. 37.

Biggiogera, M., Bottone, M.G. and Pellicciari, C. (1997), “Nuclear ribonucleoprotein-containing structures undergo severe rearrangement during spontaneous thymocyte apoptosis. A morphological study by electron microscopy”, Histochemistry and Cell Biology, Vol. 107 No. 4, pp. 331–336.

Brenner, S. (1974), “The Genetics of Caenorhabditis Elegans”, Genetics, Vol. 77 No. 1, pp. 71–94.

Burk, K. and Pasterkamp, R.J. (2019), “Disrupted neuronal trafficking in amyotrophic lateral sclerosis”, Acta Neuropathologica, available at:https://doi.org/10.1007/s00401-019-01964-7.

Cataldo, A.M., Mathews, P.M., Boiteau, A.B., Hassinger, L.C., Peterhoff, C.M., Jiang, Y., Mullaney, K., et al. (2008), “Down Syndrome Fibroblast Model of Alzheimer-Related Endosome Pathology: Accelerated Endocytosis Promotes Late Endocytic Defects”, The American Journal of Pathology, Vol. 173 No. 2, pp. 370–384.

Chang, Q. and Martin, L.J. (2016), “Voltage-gated calcium channels are abnormal in cultured spinal motoneurons in the G93A-SOD1 transgenic mouse model of ALS”, Neurobiology of Disease, Vol. 93, pp. 78–95.

de Chaumont, F., Dallongeville, S., Chenouard, N., Hervé, N., Pop, S., Provoost, T., Meas-Yedid, V., et al. (2012), “Icy: an open bioimage informatics platform for extended reproducible research”, Nature Methods, Vol. 9 No. 7, pp. 690–696.

Chou, SM. (1992), “Pathology: light microscopy of amyotrophic lateral sclerosis”, Handbook of Amyotrophic Lateral Sclerosis, pp. 133–181.

Colacurcio, D.J., Pensalfini, A., Jiang, Y. and Nixon, R.A. (2018), “Dysfunction of Autophagy and Endosomal-Lysosomal Pathways: Roles in Pathogenesis of Down Syndrome and Alzheimer’s Disease”, Free Radical Biology & Medicine, Vol. 114, pp. 40–51.

David, D.C., Ollikainen, N., Trinidad, J.C., Cary, M.P., Burlingame, A.L. and Kenyon, C. (2010), “Widespread Protein Aggregation as an Inherent Part of Aging in C. elegans”, PLOS Biology, Vol. 8 No. 8, p. e1000450.

Fogarty, M.J. (2018), “Driven to decay: Excitability and synaptic abnormalities in amyotrophic lateral sclerosis”, Brain Research Bulletin, Vol. 140, pp. 318–333.

Fogarty, M.J. (2019), “Amyotrophic lateral sclerosis as a synaptopathy”, Neural Regeneration Research, Vol. 14 No. 2, pp. 189–192.

Gal, J., Zhang, J., Kwinter, D.M., Zhai, J., Jia, H., Jia, J. and Zhu, H. (2011), “Nuclear localization sequence of FUS and induction of stress granules by ALS mutants”, Neurobiology of Aging, Vol. 32 No. 12, pp. 2323.e27-2323.e40.

Gao, S. and Zhen, M. (2011), “Action potentials drive body wall muscle contractions in Caenorhabditis elegans”, Proceedings of the National Academy of Sciences, Vol. 108 No. 6, pp. 2557–2562.

Guo, W., Naujock, M., Fumagalli, L., Vandoorne, T., Baatsen, P., Boon, R., Ordovás, L., et al. (2017), “HDAC6 inhibition reverses axonal transport defects in motor neurons derived from FUS-ALS patients”, Nature Communications, Vol. 8 No. 1, pp. 1–15.

Hardiman, O., Al-Chalabi, A., Chio, A., Corr, E.M., Logroscino, G., Robberecht, W., Shaw, P.J., et al. (2017), “Amyotrophic lateral sclerosis”, Nature Reviews Disease Primers, Vol. 3, p. 17071.

Hicks, G.G., Singh, N., Nashabi, A., Mai, S., Bozek, G., Klewes, L., Arapovic, D., et al. (2000), “Fus deficiency in mice results in defective B-lymphocyte development and activation, high levels of chromosomal instability and perinatal death”, Nature Genetics, Vol. 24 No. 2, p. 175.

Holt, C.E. and Schuman, E.M. (2013), “The Central Dogma Decentralized: New Perspectives on RNA Function and Local Translation in Neurons”, Neuron, Vol. 80 No. 3, pp. 648–657.

Huang, C., Wagner-Valladolid, S., Stephens, A.D., Jung, R., Poudel, C., Sinnige, T., Lechler, M.C., et al. (2019), “Intrinsically aggregation-prone proteins form amyloid-like aggregates and contribute to tissue aging in Caenorhabditis elegans”, edited by Kuriyan, J.ELife, Vol. 8, p. e43059.

Jokhi, V., Ashley, J., Nunnari, J., Noma, A., Ito, N., Wakabayashi-Ito, N., Moore, M.J., et al. (2013), “Torsin Mediates Primary Envelopment of Large Ribonucleoprotein Granules at the Nuclear Envelope”, Cell Reports, Vol. 3 No. 4, pp. 988–995.

Jung, H., Gkogkas, C.G., Sonenberg, N. and Holt, C.E. (2014), “Remote Control of Gene Function by Local Translation”, Cell, Vol. 157 No. 1, pp. 26–40.

Kaltdorf, K.V., Schulze, K., Helmprobst, F., Kollmannsberger, P., Dandekar, T. and Stigloher, C. (2017), “FIJI Macro 3D ART VeSElecT: 3D Automated Reconstruction Tool for Vesicle Structures of Electron Tomograms”, PLOS Computational Biology, Vol. 13 No. 1, p. e1005317.

Kaltdorf, K.V., Theiss, M., Markert, S.M., Zhen, M., Dandekar, T., Stigloher, C. and Kollmannsberger, P. (2018), “Automated classification of synaptic vesicles in electron tomograms of C. elegans using machine learning”, PLOS ONE, Vol. 13 No. 10, p. e0205348.

Kino, Y., Washizu, C., Aquilanti, E., Okuno, M., Kurosawa, M., Yamada, M., Doi, H., et al. (2011), “Intracellular localization and splicing regulation of FUS/TLS are variably affected by amyotrophic lateral sclerosis-linked mutations”, Nucleic Acids Research, Vol. 39 No. 7, pp. 2781– 2798.

Kong, L., Wang, X., Choe, D.W., Polley, M., Burnett, B.G., Bosch-Marcé, M., Griffin, J.W., et al. (2009), “Impaired Synaptic Vesicle Release and Immaturity of Neuromuscular Junctions in Spinal Muscular Atrophy Mice”, Journal of Neuroscience, Vol. 29 No. 3, pp. 842–851.

Kremer, J.R., Mastronarde, D.N. and McIntosh, J.R. (1996), “Computer Visualization of Three-Dimensional Image Data Using IMOD”, Journal of Structural Biology, Vol. 116 No. 1, pp. 71– 76.

Kwiatkowski, T.J., Bosco, D.A., LeClerc, A.L., Tamrazian, E., Vanderburg, C.R., Russ, C., Davis, A., et al. (2009), “Mutations in the FUS/TLS Gene on Chromosome 16 Cause Familial Amyotrophic Lateral Sclerosis”, Science, Vol. 323 No. 5918, pp. 1205–1208.

Lagier-Tourenne, C., Polymenidou, M. and Cleveland, D.W. (2010), “TDP-43 and FUS/TLS: emerging roles in RNA processing and neurodegeneration”, Human Molecular Genetics, Vol. 19 No. R1, pp. R46–R64.

Liu, M.-L., Zang, T. and Zhang, C.-L. (2016), “Direct Lineage Reprogramming Reveals Disease-Specific Phenotypes of Motor Neurons from Human ALS Patients”, Cell Reports, Vol. 14 No. 1, pp. 115– 128.

López-Erauskin, J., Tadokoro, T., Baughn, M.W., Myers, B., McAlonis-Downes, M., Chillon-Marinas, C., Asiaban, J.N., et al. (2018), “ALS/FTD-Linked Mutation in FUS Suppresses Intra-axonal Protein Synthesis and Drives Disease Without Nuclear Loss-of-Function of FUS”, Neuron, Vol. 100 No. 4, pp. 816–830.e7.

Lorenzo-Betancor, O., Ogaki, K., Soto-Ortolaza, A., Labbé, C., Vilariño-Güell, C., Rajput, A., Rajput, A.H., et al. (2014), “Analysis of Nuclear Export Sequence Regions of FUS-Related RNA-Binding Proteins in Essential Tremor”, PLOS ONE, Vol. 9 No. 11, p. e111989.

Markert, S.M., Bauer, V., Muenz, T.S., Jones, N.G., Helmprobst, F., Britz, S., Sauer, M., et al. (2017), “Chapter 2 - 3D subcellular localization with superresolution array tomography on ultrathin sections of various species”, in Müller-Reichert, T. and Verkade, P. (Eds.), Methods in Cell Biology, Vol. 140, Academic Press, pp. 21–47.

Markert, S.M., Britz, S., Proppert, S., Lang, M., Witvliet, D., Mulcahy, B., Sauer, M., et al. (2016), “Filling the gap: adding super-resolution to array tomography for correlated ultrastructural and molecular identification of electrical synapses at the C. elegans connectome”, Neurophotonics, Vol. 3 No. 4, pp. 041802–041802.

Martínez-Silva, M. de L., Imhoff-Manuel, R.D., Sharma, A., Heckman, C., Shneider, N.A., Roselli, F., Zytnicki, D., et al. (2018), “Hypoexcitability precedes denervation in the large fast-contracting motor units in two unrelated mouse models of ALS”, edited by Rubin, L.L.ELife, Vol. 7, p. e30955.

Mastronarde, D.N. (2005), “Automated electron microscope tomography using robust prediction of specimen movements”, Journal of Structural Biology, Vol. 152 No. 1, pp. 36–51.

Meléndez, A., Tallóczy, Z., Seaman, M., Eskelinen, E.-L., Hall, D.H. and Levine, B. (2003), “Autophagy Genes Are Essential for Dauer Development and Life-Span Extension in C. elegans”, Science, Vol. 301 No. 5638, pp. 1387–1391.

Murakami, T., Qamar, S., Lin, J.Q., Schierle, G.S.K., Rees, E., Miyashita, A., Costa, A.R., et al. (2015), “ALS/FTD Mutation-Induced Phase Transition of FUS Liquid Droplets and Reversible Hydrogels into Irreversible Hydrogels Impairs RNP Granule Function”, Neuron, Vol. 88 No. 4, pp. 678–690.

Murakami, T., Yang, S.-P., Xie, L., Kawano, T., Fu, D., Mukai, A., Bohm, C., et al. (2012), “ALS mutations in FUS cause neuronal dysfunction and death in Caenorhabditis elegans by a dominant gain-of-function mechanism”, Human Molecular Genetics, Vol. 21 No. 1, pp. 1–9.

Naujock, M., Stanslowsky, N., Bufler, S., Naumann, M., Reinhardt, P., Sterneckert, J., Kefalakes, E., et al. (2016), “4-Aminopyridine Induced Activity Rescues Hypoexcitable Motor Neurons from Amyotrophic Lateral Sclerosis Patient-Derived Induced Pluripotent Stem Cells”, STEM CELLS, Vol. 34 No. 6, pp. 1563–1575.

Paul-Gilloteaux, P., Heiligenstein, X., Belle, M., Domart, M.-C., Larijani, B., Collinson, L., Raposo, G., et al. (2017), “eC-CLEM: flexible multidimensional registration software for correlative microscopies”, Nature Methods, Vol. 14 No. 2, pp. 102–103.

Pham-Gia, T. and Hung, T.L. (2001), “The mean and median absolute deviations”, Mathematical and Computer Modelling, Vol. 34 No. 7, pp. 921–936.

Picchiarelli, G., Demestre, M., Zuko, A., Been, M., Higelin, J., Dieterlé, S., Goy, M.A., et al. (2019), “FUS-mediated regulation of acetylcholine receptor transcription at neuromuscular junctions is compromised in amyotrophic lateral sclerosis.”, Nature Neuroscience, Vol. 22 No. 11, pp. 1793– 1805.

Ratti, A. and Buratti, E. (2016), “Physiological functions and pathobiology of TDP-43 and FUS/TLS proteins”, Journal of Neurochemistry, Vol. 138 No. S1, pp. 95–111.

Reynolds, E.S. (1963), “The use of lead citrate at high pH as an electron-opaque stain in electron microscopy”, The Journal of Cell Biology, Vol. 17 No. 1, pp. 208–212.

Richmond, J.E. and Jorgensen, E.M. (1999), “One GABA and two acetylcholine receptors function at the C. elegans neuromuscular junction”, Nature Neuroscience, Vol. 2 No. 9, pp. 791–797.

Ruiz, R., Casañas, J.J., Torres-Benito, L., Cano, R. and Tabares, L. (2010), “Altered Intracellular Ca2+ Homeostasis in Nerve Terminals of Severe Spinal Muscular Atrophy Mice”, Journal of Neuroscience, Vol. 30 No. 3, pp. 849–857.

Schindelin, J., Arganda-Carreras, I., Frise, E., Kaynig, V., Longair, M., Pietzsch, T., Preibisch, S., et al. (2012), “Fiji: an open-source platform for biological-image analysis”, Nature Methods, Vol. 9 No. 7, pp. 676–682.

Souquere, S., Mollet, S., Kress, M., Dautry, F., Pierron, G. and Weil, D. (2009), “Unravelling the ultrastructure of stress granules and associated P-bodies in human cells”, Journal of Cell Science, Vol. 122 No. 20, pp. 3619–3626.

Stiernagle, T. (2006), “Maintenance of C. elegans”, WormBook: The Online Review of C. Elegans Biology, pp. 1–11.

Stigloher, C., Zhan, H., Zhen, M., Richmond, J. and Bessereau, J.-L. (2011), “The presynaptic dense projection of the Caenorhabiditis elegans cholinergic neuromuscular junction localizes synaptic vesicles at the active zone through SYD-2/liprin and UNC-10/RIM-dependent interactions”, The Journal of Neuroscience : The Official Journal of the Society for Neuroscience, Vol. 31 No. 12, pp. 4388–4396.

Therrien, M., Rouleau, G.A., Dion, P.A. and Parker, J.A. (2016), “FET proteins regulate lifespan and neuronal integrity”, Scientific Reports, Vol. 6, p. 25159.

Vance, C., Rogelj, B., Hortobágyi, T., Vos, K.J.D., Nishimura, A.L., Sreedharan, J., Hu, X., et al. (2009), “Mutations in FUS, an RNA Processing Protein, Cause Familial Amyotrophic Lateral Sclerosis Type 6”, Science, Vol. 323 No. 5918, pp. 1208–1211.

Vance, C., Scotter, E.L., Nishimura, A.L., Troakes, C., Mitchell, J.C., Kathe, C., Urwin, H., et al. (2013), “ALS mutant FUS disrupts nuclear localization and sequesters wild-type FUS within cytoplasmic stress granules”, Human Molecular Genetics, Vol. 22 No. 13, pp. 2676–2688.

Walther, D.M., Kasturi, P., Zheng, M., Pinkert, S., Vecchi, G., Ciryam, P., Morimoto, R.I., et al. (2015), “Widespread Proteome Remodeling and Aggregation in Aging C. elegans”, Cell, Vol. 161 No. 4, pp. 919–932.

Wang, H., Guo, W., Mitra, J., Hegde, P.M., Vandoorne, T., Eckelmann, B.J., Mitra, S., et al. (2018), “Mutant FUS causes DNA ligation defects to inhibit oxidative damage repair in Amyotrophic Lateral Sclerosis”, Nature Communications, Vol. 9 No. 1, p. 3683.

Watanabe, S., Liu, Q., Davis, M.W., Hollopeter, G., Thomas, N., Jorgensen, N.B. and Jorgensen, E.M. (2013), “Ultrafast endocytosis at Caenorhabditis elegans neuromuscular junctions”, ELife, Vol. 2013 No. 2, p. e00723.

Watanabe, S., Rost, B.R., Camacho-Pérez, M., Davis, M.W., Söhl-Kielczynski, B., Rosenmund, C. and Jorgensen, E.M. (2013), “Ultrafast endocytosis at mouse hippocampal synapses”, Nature, Vol. 504 No. 7479, pp. 242–247.

Watanabe, S., Trimbuch, T., Camacho-Pérez, M., Rost, B.R., Brokowski, B., Söhl-Kielczynski, B., Felies, A., et al. (2014), “Clathrin regenerates synaptic vesicles from endosomes”, Nature, Vol. 515 No. 7526, pp. 228–233.

Weimer, RobbyM. (2006), “Preservation of C. elegans Tissue Via High-Pressure Freezing and Freeze-Substitution for Ultrastructural Analysis and Immunocytochemistry”, in Strange, K. (Ed.), C. Elegans, Humana Press, pp. 203–221.

Yan, D., Wu, Z., Chisholm, A.D. and Jin, Y. (2009), “The DLK-1 Kinase Promotes mRNA Stability and Local Translation in C. elegans Synapses and Axon Regeneration”, Cell, Vol. 138 No. 5, pp. 1005–1018.

